# Activities of aqueous extract of *Tamarindus indica* on phenylhydrazine-induced hematopathological changes in anemic male wistar rats

**DOI:** 10.64898/2026.01.30.702729

**Authors:** Cherop Patrick Stephen, Edgar Fernandez Mario, Dare Samuel Sunday

**Affiliations:** Department of Anatomy, Faculty of Biomedical Sciences, Kampala International University, Western Campus, Uganda; Department of Anatomy, School of Medicine, Kabale University, Uganda

**Author notes:** **Corresponding Author**: Samuel Sunday Dare, School of Medicine, Kabale University, P.O. Box 317, Kabale, Uganda. These authors contributed equally to this work (Cherop Patrick Stephen, Dare Samuel Sunday).

**Keywords:** PHZ, *T. indica*, LDH, RBCs, hemolytic anemia, reticulocytes, bone marrow, hematopoiesis

## Abstract

**Introduction:** Drug Induced Hemolytic Anemia (DIHA), following exposure to hematopathologically profound molecules, presents with variable clinical syndromes, misinterpreted serological results, misdiagnosis, challenging and controversial treatment; and no specific antihemolytic agent. Its treatment could be enhanced by use of natural molecules in some medicinal plants. Therefore, this study is aimed to determination the activities of aqueous extract of *T. indica* on PHZ-induced hematopathological changes in anemic male *Wistar rats*.

**Materials and Methods:** 60mg/kg of Phenylhydrazine (PHZ) was administered for 2 days to induce hemolytic anemia intraperitoneally. 30 male *Wistar rats* were randomly divided into 5 groups, each with 6 rats. G_1_-untreated. Anemic rats were divided into G_2_- G_5_. G_2_-untreated, G_3_-treated with 1mL Ferro B syrup, G_4_ and G_5_ treated with 400mg/kg and 800mg/kg of *T. indica* pulp extract respectively. Test drug and extract were orally administered daily for 7 and 14 days respectively. Cases in G_2_ - G_5_ were sacrificed under light ether anesthesia on days 9 and 16 post-therapy, G_1_ at the end of the experimental period. Blood collected via cardiac puncture were subjected for Red Blood Cells (RBC) histopathology, serum Lactate Dehydrogenase (LDH), and reticulocyte counts. The femur was harvested for bone marrow Histopathology.

**Results:** PHZ induced hemolytic anemia marked by profound serum LDH elevation & reticulocytosis, marked RBC morphological distortions & bone marrow degenerative changes suggestive of marrow fibrosis & suppression. Marrow regeneration marked by hypercellularity & decreased adipocytes were evident of hematopoiesis induced by the 2 weeks test therapies; significant moderate populations of normal mature peripheral RBCs, serum LDH & reticulocyte % reduction were typical; consistent with significant recovery from the acute hemolytic episode.

**Conclusion:** *T. indica* fruit pulp extract effectively stimulated hematopoiesis in response to drug induced hemolytic effect on the hematopathologic parameters, with significant improvement from hemolytic anemia.

## Introduction

Hemolytic anemia, known to be associated with acutely or chronically occurring hemolysis, is precipitated by known or unknown triggers. Hemolysis releases hemoglobin and other intracellular components into the plasma [1], shortens RBC life span, directly or indirectly effecting hematopoietic organs, resulting in clinical vs. anatomic pathological effects like: reticulocytosis with correlative changes in RBC indices; increased splenic and bone marrow (BM) erythropoiesis, splenomegally, splenic congestion and pigment [2].

Drug induced hemolysis (DIH) is linked to immunological and metabolic phenomena. Autoantibodies react with target cells in presence or absence of the drug causing antibody mediated extravascular hemolysis; drug dependent antibodies bind to RBCs only in the presence of sensitizing drug leading to complement-mediated intravascular hemolysis [3]. However, some drugs have capability of stimulating the production of both types of antibodies [4].

Immunologically induced hemolytic mechanisms including hapten induced - active drug molecular components attach to the RBC membrane stimulating IgG antibody production, complex immune mechanism-induces immunoglobulin (Ig)M antibody production where the drug antibody complex binds to the RBC membrane initiating complement activation with resultant intravascular hemolysis and anti-RBC IgG antibody produced by induction of the autoantibody causes extravascular hemolysis by unknown mechanism have been proposed [5].

The metabolic phenomenon is associated with oxidative stress induced by the drug strongly linked to RBC enzymopathies, especially Glucose-6-Phosphate Dehydrogenase (G6PD) characterized by Nicotinamide Adenine Dinucleotide Phosphate (NADPH) production leading to altered state of Glutathione and consequential defective antioxidant function, creating an advantage for oxidant drugs that oxidize the Glutathione groups of Hemoglobin (Hb), removing them prematurely from circulation, and causing hemolysis [6]. The free Hb exerts direct cytotoxic, inflammatory and pro-oxidants effects [7], induce membrane lipid peroxidation [8], generate Reactive Oxygen Species (ROS), chemokines, cytokines and GFs [9] exerting additional hematopoietic demand in the BM, and enhanced splenic erythropoiesis [10].

Drugs may interfere with the hematopoietic stem cells’ DNA replication resulting in genetic alterations and/or damage that may either directly impair the stem cell’s proliferation and differentiation capacity or induce neoantigen expression in the stem cell and its progeny indirectly, triggering immune destruction via recruitment of cytotoxic T lymphocytes [11]. PHZ induces hemolytic anemia via oxidative degradation of some RBC membrane proteins impairing its deformability, destabilizing the Hb by replacing them with phenyl radicals, and multiple other cellular changes [12].

Apparently, there are no available specific anti-hemolytic drugs. Those used may control DIH via unknown mechanisms which are at the same time directed against the immune system, creating a clinician-patient dilemma. However, included among the treatment options are traditional treatments [13] which are believed to be useful in strengthening the hematopoietic and immune system, and treatment of hematological disorders [14], supported by the fact that they contain remarkable physiologically active phytochemicals [15].

*T. indica* is used in West Nile region of Uganda as a preventive measure during epidemic diseases’ outbreak [16], for treatment of meningitis and other ailments [17]. Its phytochemical components have been found to stabilize the RBC membrane, preventing their damage [18]. This plants’ ability to restore hemato-biochemical parameters and maintain oxidative balance in rats, and other mammals like cattle and rabbits has been reported [19, 20, 21, 22]. This potential has been extended to many chronic diseases [23], pointing out the possibility of its usefulness in the management of hemolytic anemia secondary to drugs or other causes. Despite the ameliorative effects evaluated in experimental animals, no report on its activities on drug induced hemolytic anemia in *Wistar rats* have been fully documented, particularly in Uganda, prompting this study to evaluate its activities on PHZ-induced hemolytic anemia in *Wistar rats*, addressing the RBC histomorphological changes and bone marrow cyto-architectural changes; and some key indirect hemolytic markers.

## Materials and methods

### Herbal medicine

*T. indica* pods were procured from the local markets in the Eastern Districts of Kumi, Mayuge, and Jinja respectively because of their geographical distribution [17], and were authenticated from the Department of Botany. The aqueous fruit pulp extract of the *T. indica* pulp was prepared and used as the herbal medicine.

### Preparation of the aqueous extract

The pods were washed with clean water and allowed to dry for 1 hour to maintain constant moisture content of the shell. Pod shells were manually depulped with decomposed and damaged pulps discarded. The selected pulps were then placed in air-tight polythene bag to keep them fresh. The aqueous extract was prepared by the method described by Rosenthaler [24] and Azwanida [25]. 10% aqueous pulp extract was prepared from 500g of the fresh pulp which was blended using an electric blender, diluted with distilled water in the ratio of 1:10, and placed in a conical flask tightly stoppered with Aluminum foil. This conical flask was then placed in an orbital shaker at 125rpm for 3 days, at room temperature. These contents were then filtered through muslin cloth. The residue left in the flask was rinsed with little quantity of distilled water and filtered through the muslin cloth. The filtrate obtained was finally filtered through Whatman filter paper No. 1 to obtain final filtrate, which was evaporated in an oven at 40°C for 3 days. 100g of the final extract was obtained and stored at at 4°C in a refrigerator for further use. 25g of the extract was then reconstituted with 250mls of distilled water to a concentration of 100mg/ml, which was administered to the rats at respective doses.

### Preparation of PHZ

PHZ powder (Cat. No. 11,471-5, Sigma-Aldrich, USA) was used. 1g of the powder was reconstituted with 50ml of distilled water to a concentration of 20mg/ml. A dosage of 60mg/kg body weight was used against the 20mg/ml concentration to get the desired volume to administer to each weighed rat.

### Experimental animals

Thirty (30) male Wistar rats weighing between 130-180g were procured from the animal breeding house of Kampala International University-Western Campus. They were kept in cages under a 12-hour light and 12-hour dark period with each cage lined with rat bedding material that was changed every alternate day. The animals were fed on standard rat pellet diet and clean water and were allowed to acclimatize to the environment prior to experimentation.

### Animal grouping and treatment

The animals were divided into 5 groups (G_1_, G_2_, G_3_, G_4_, and G_5)_, each consisting of 6 rats (n = 6) with each rat given a specific unique identification mark. G_1_ served as normal control and received adequate amount of clean water per day. G_2_ served as untreated and received 60mg/kg of I.P PHZ for 2 days. G_3_ served as experimental group I (anemic) and received 1mL of stand hematinic drug (Ferro B-complex syrup) *per os* daily. G_4_ served as experimental group II (anemic) and received 400mg/kg of *T. indica* aqueous fruit pulp extract *per os* whereas G_5_ served as experimental group III (anemic) and received 800mg/kg of *T. indica* aqueous fruit pulp extract *per os*.

### Experimental design

Acclimatized healthy male Wistar rats were induced with hemolytic anemia using PHZ. Hemolytic anemia was first induced in 5 separate rats, sacrificed to obtain 5 ml of cardiac blood which was assessed for confirmation of hemolytic anemia.

On day 0, hemolytic anemia was induced in 24 rats (G_2_, G_3_, G_4_, and G_5_) by administration of 60mg/kg of PHZ intraperitoneally daily for two days. The treatment of the anemic rats then commenced on day three, with the extract of *T. indica* administered orally for 14 days. On days 9 & 16, 3 rats per group were sacrificed under light ether anaesthesia [26], and 5 ml of cardiac blood was obtained by cardiac puncture. 2 ml was placed in EDTA-coated vacutainers while 3 ml was placed in red topped vacutainers. The blood in the red topped vacutainers was centrifuged at 2000 rpm for 15 min to separate the serum from the clot. The serum was pipetted into labeled cuvettes for serum LDH analysis [27], while blood in the EDTA-coated vacutainers were analysed for reticulocytes and RBC morphological changes [27]. The bone marrow was also harvested for histopathology study.

### Hemato-biochemical assessment

#### RBC morphology

Thin blood smears were prepared following the procedure described by Gulati *et al*. [28]. Approximately 25µl of blood sample was placed and spread using a glass spreader on a microscopic glass slide; air dried, fixed with absolute methanol and again allowed to dry for 4 minutes. Giemsa stain was added on the glass slide, washed with de-ionized water, then allowed to dry. The slides were mounted on the stage and viewed under x100 objective lens and pictures taken using camera fitted microscope. Morphological changes detected in the treated groups were compared with the respective control groups (Figure 6-12).

#### LDH

This was determined according to the method described by Kumari *et al*. [29]. 20 ml of the working solution was prepared and 1 ml pipetted into each cuvette. 20μl of the rat serum samples was added to each of the cuvettes. 1ml of the LDH assay buffer was then added to each cuvette. The contents were well mixed using a pipette, incubated at 37°C for 30 seconds, and the initial absorbance at 340 nm was read after 1 minute. The stop watch was started simultaneously and further readings of the absorbance were taken at exactly 1, 2, and 3 minutes. The change in absorbance per minute (ΔA/min) was determined and the LDH activity calculated using the formulae: U/l = 8095 × ΔA 340 nm/min, where 8095 is the Extinction coefficient.

#### Reticulocytes

These were determined according to the method described by Bain *et al*. [30]. 1.0 g of new methylene blue was dissolved in 100 ml of 3% trisodium citrate-saline solution (30g of sodium citrate in 11 of saline). The mixture was filtered once into a beaker. 2 drops of the dye solution was pipette using a plastic Pasteur pipette into a 75 × 10 mm plastic tube. Adequate volume of the rat’s EDTA-anticoagulated blood was added to the dye solution and mixed. This mixture was then incubated at 37°C for 15-20 min. The RBCs were then suspended by gentle mixing and two films for each rat were made on glass slides, gently well spread and left to dry, and were examined without fixing or counterstaining at x100 oil immersion objective. The reticulocytes were counted together with the total RBCs per field in 10 fields. The reticulocyte percentages were determined as follows:

Number of reticulocytes in *n* fields = *x* Average number of red cells per field = *y* Total number of red cells in *n* fields = *n* × *y* Reticulocyte percentage = [*x* ÷ (*n* × y)] × 100

### Bone marrow preparation for histomorphological analysis

The bone marrow was harvested from the femur after the rats were sacrificed. The dissected femur bone was removed and excess muscle and fat trimmed off. The bone was split longitudinally to expose the marrow. A fine brush was used to gently extract a small plug of the marrow which was smeared on the glass slides, air-dried, fixed with methanol and stained using the Wright-Giemsa which highlights the nuclear and cytoplasmic details [31].

Another femur was removed, fixed immediately in 10% neutral buffered formalin for 24-48 hours, decalcified with formic acid-formalin solution and then processed for paraffin embedded histology. Sections were cut out at 6 µm thickness and processed according to Comanescu *et al*.[32], stained with H & E (Hematoxylin and Eosin) followed by examination under camera fitted microscope, and pictures taken. The stained slide films were examined under camera fitted binocular microscope at medium power (x20) and high power (x40) magnifications. Histomorphological analysis of bone marrow cyto-architecture, assessment of cellularity and other related degenerative and regenerative features were done (Figure 12 & 13).

### Data analysis

Data was stored in MS-Excel and statistical analysis carried using SPSS where One-Way ANOVA was employed to determine the statistical significance set at (p<0.05) at 95% confidence interval. Tukey’s post hoc analysis was used to test for the differences in the means between the groups.

### Ethical statement

The experimental protocol, having met the National Institute of Health guidelines for the care and use of the laboratory animals, was approved by the Institutional Rights Research and Ethics Committee of Kampala International University (IREC-KIU). The experiment was conducted in compliance with the National Institute of Health Guidelines for Care and Use of Laboratory Animals.

## Results

Five fatalities were registered. Statistical analysis of the data for the rest of the subjects was presented (Table 1).

**Table 1:** Changes in the serum LDH and Reticulocyte percentage (retic. %) of the rats.

| Experimental groups |  |  |  |  |  |  |  |
| --- | --- | --- | --- | --- | --- | --- | --- |
| Parameter | Rx period | G1 | G2 | G3 | G4 | G5 | P value |
| LDH(IU/L) | I | 200.5±26.13 | 360.0±28.28 | 318.0±14.14 | 320.0±5.65 | 316.0±5.65 | 0.003 |
|  | II | 137.5±17.67 | 302.0±2.82 | 289.0±2.82 | 279.0±5.65 | 261.0±8.48 | 0.0001 |
| Retic. % | I | 2.55±0.35 | 16.15±0.63 | 15.1±0.28 | 13.9±0.84 | 12.4±0.84 | 0.0001 |
|  | II | 1.75±0.21 | 11.45±0.77 | 10.5±1.13 | 10.0±1.69 | 8.97±1.55 | 0.003 |

Reticulocytosis is marked by increased reticulocyte percentage >5% beyond the normal range (1%-5%). The LDH normal range: 120-240 IU/L. Values are in Mean±SD; G1= normal control; G2=anemic untreated; G3=anemic treated with Ferro B syrup; G4=anemic treated with 400mg/kg of *T.indica* fruit pulp extract; G5= anemic treated with 800mg/kg of *T.indica* fruit pulp extract; I= 7 days of treatment; II=14 days of treatment. One Way ANOVA and Tukey's HSD post hoc were used, P value considered significant if $p < 0.05$ .

### Reticulocyte percentage

The reticulocyte % remained considerably high in all the anemic rats treated with ferro B syrup and the *T.indica* fruit pulp extract, but was significantly higher (16.15±0.63, *p***<**0.05) in the untreated anemic rats, one week following PHZ-induced acute hemolytic event. A slightly significant decrease in the reticulocyte percentage was seen in all the rats, untreated and treated, after 14 days of treatment. This decrease was more marked (8.97±1.55, *p***<**0.05) in the rats treated with 800mg/kg of the *T.indica* fruit pulp extract (Table 1, figure 1 & 2).

**Figure 1:**
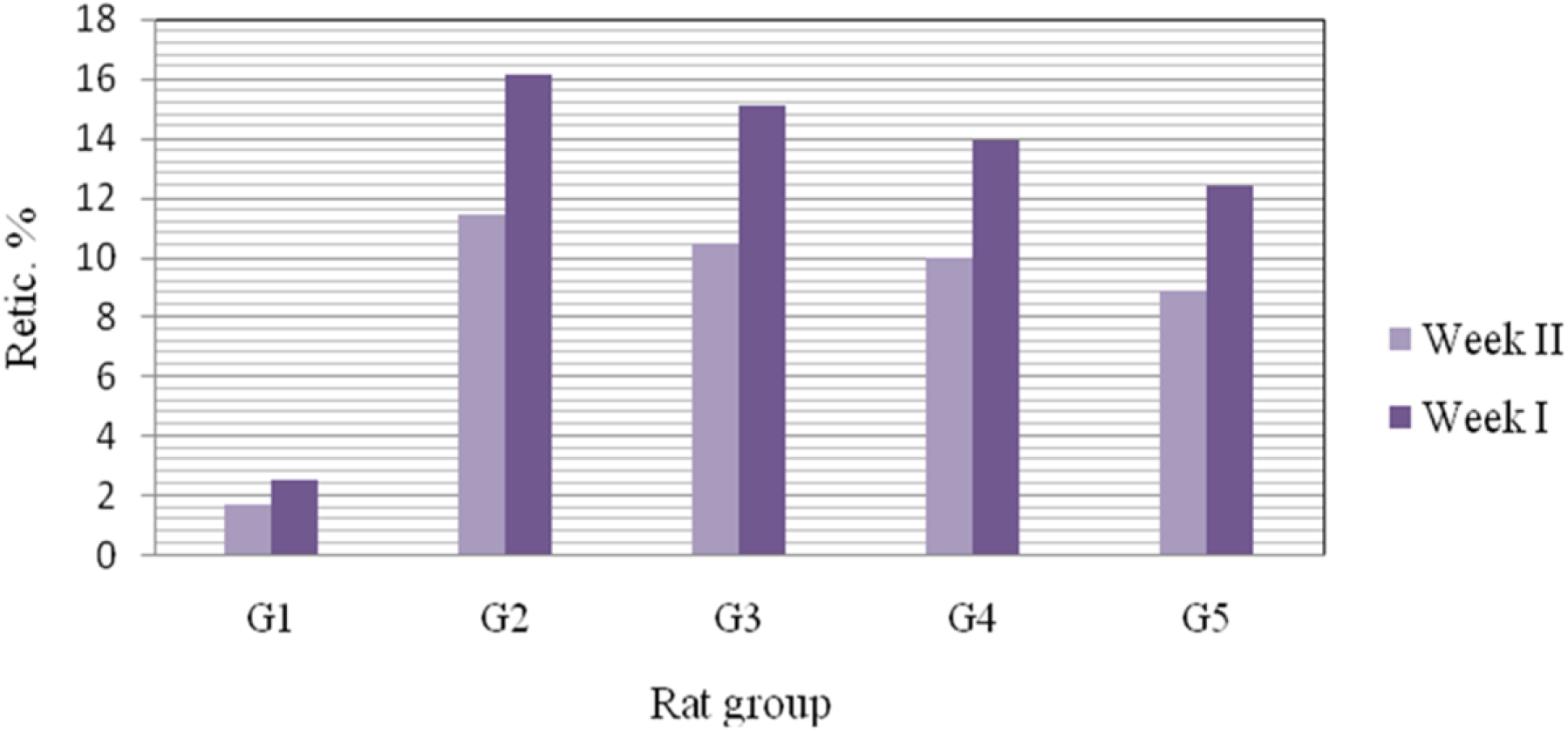
Mean±SD of the Reticulocyte percentages of the rat groups after the experimental periods. Marked reticulocytosis was seen among G2 rats one week post PHZ-induction.

**Figure 2:**
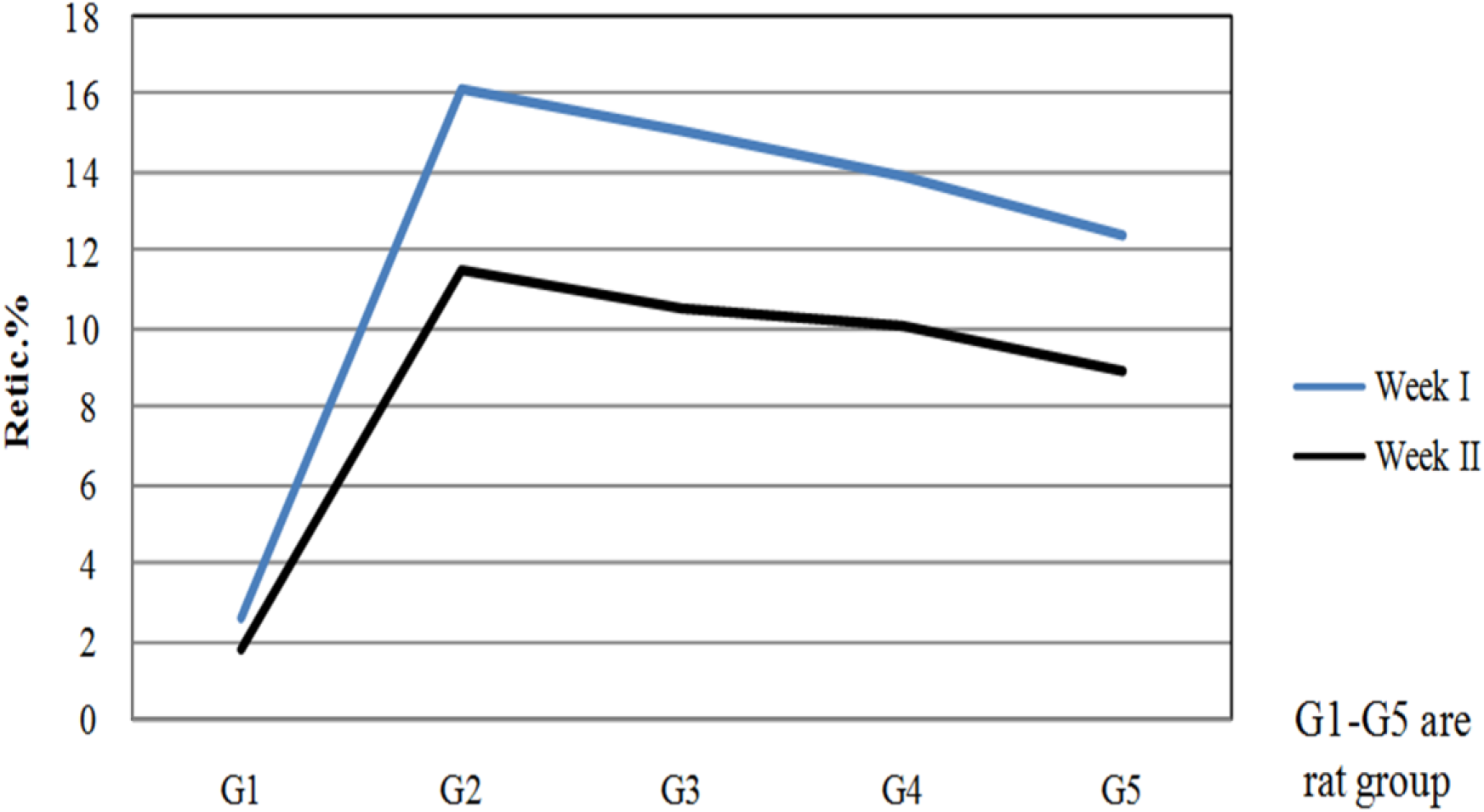
Mean±SD of the Reticulocyte percentages of the rat groups after the experimental periods

### Serum LDH levels

Similarly, it was observed that reticulocyte percentage correlated with the serum LDH level. Though the serum LDH for the untreated rats remained significantly high (360.0±28.28, *p***<**0.05) after one week, a spontaneous significant drop (302.0±2.82, *p***<**0.05) was seen by the end of the second week, which cut across all the treated groups. However, the drops in the serum LDH of the treated rats by the end of the first week were not statistically significant (Table 2, figure 3 & 4).

**Figure 3:**
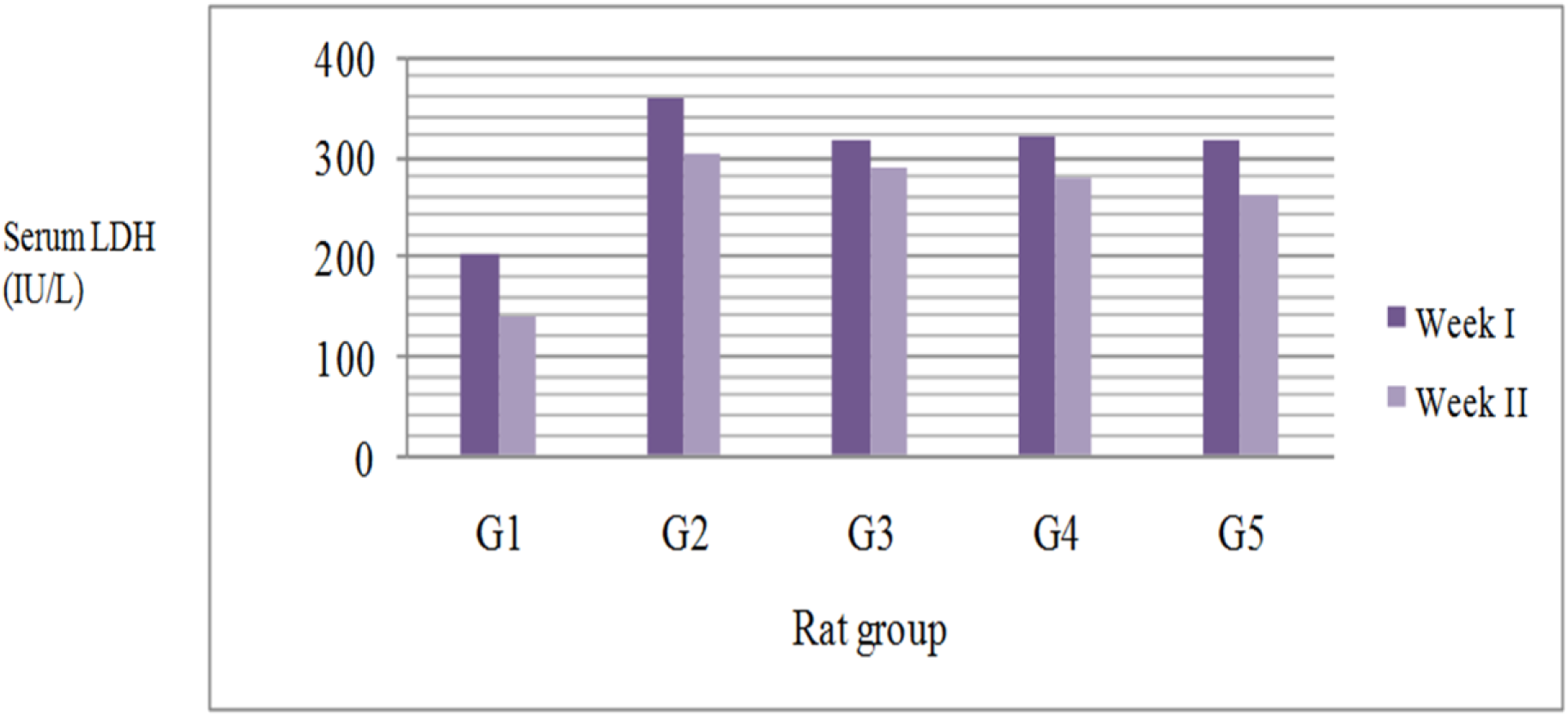
Mean±SD of the serum LDH of the rat groups after the experimental periods. After one week of treatment, G2 had nearly twice the serum LDH compared to that of G1; G3 to G4 had a sustained serum LDH of about 16% (58 IU/L). After two weeks of treatment, the serum LDH level of G2 remained significantly high, with a mean difference of about 16% (58 IU/L).

**Figure 4:**
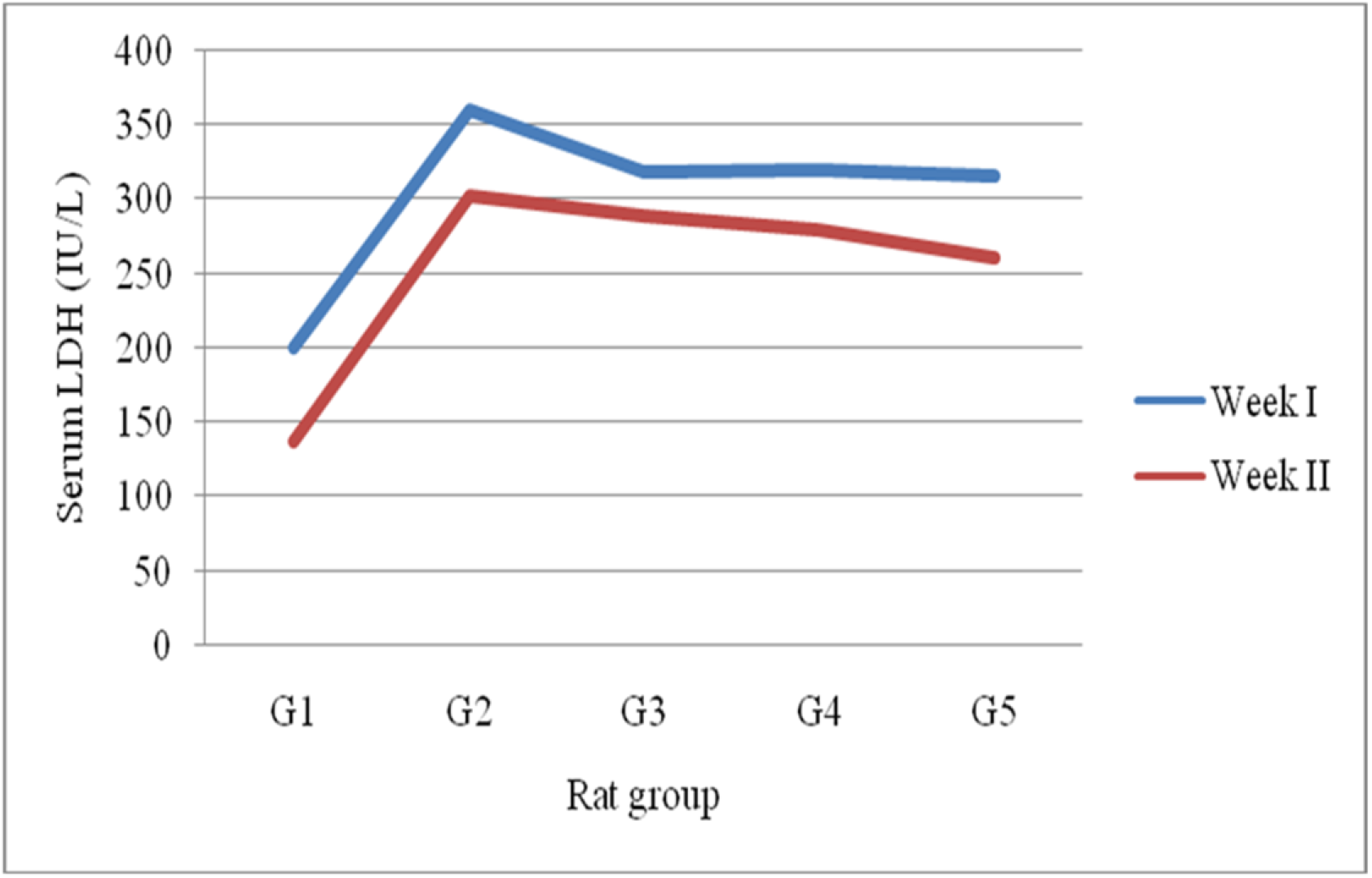
Mean±SD of the serum LDH of the rat groups after the experimental periods.

### High power RBC histomorphology results pre and post-therapy (extract, ferro B syrup and PHZ)

#### Normal control group

For the normal control, the erythrocytes were of normal size and showed characteristic central pallor. The rouleaux seen were of no diagnostic significance (figure 5).

**Figure 5:**
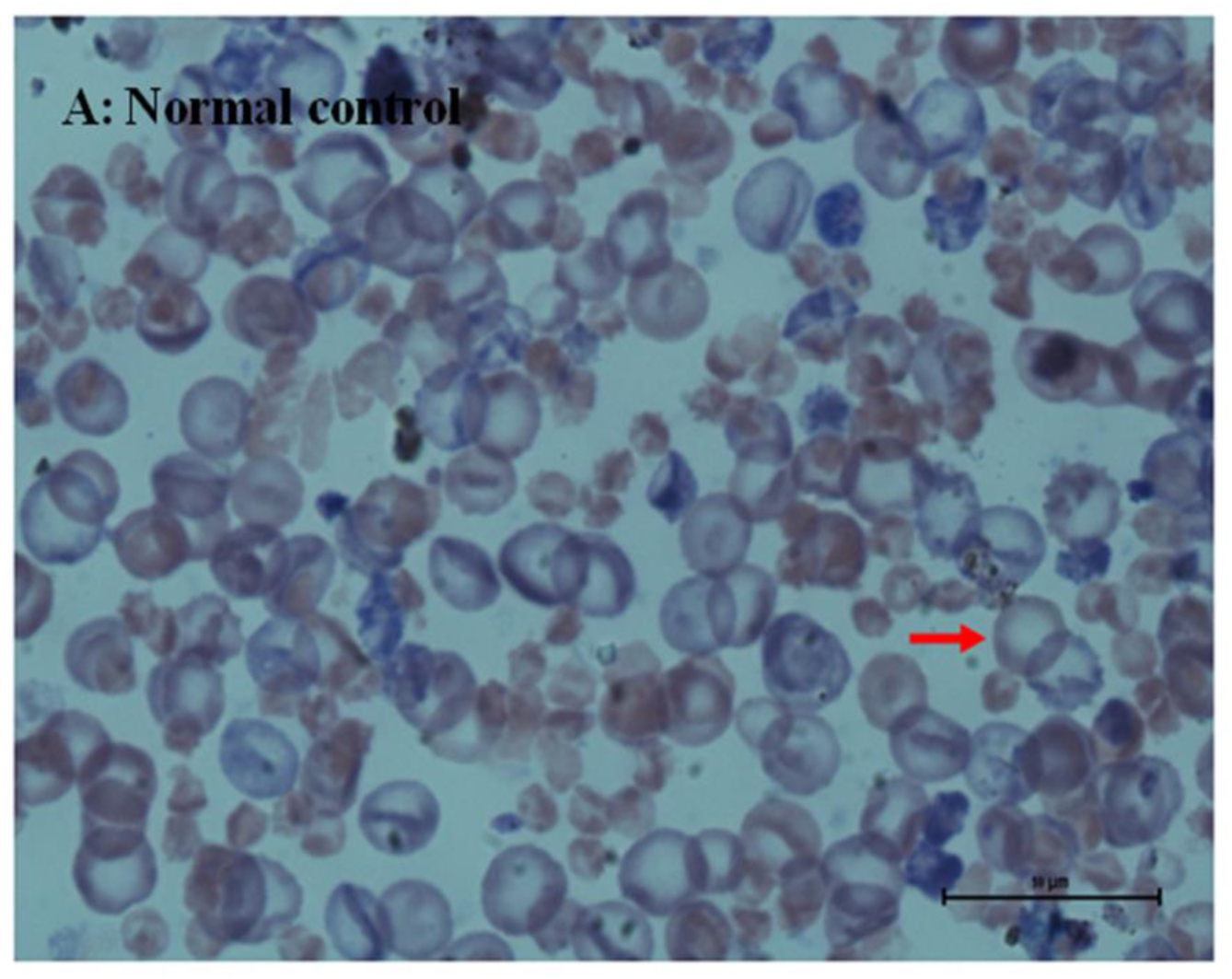
Blood smear, RBCs, Giemsa, x 100. The bold red arrow points to erythrocytes.

#### Anemic-untreated group

After one week post-PHZ induced acute hemolytic event, there was Spherocytic predominance, with bite cells, blister cells and many acanthocytes/spur cells. Few echinocytes/burr cells and tear drop cells were also present (Figure 6). Few normal sized & well hemoglobinised RBCs, faint blue cells, bite cells and moderately increased spherocytes of varied sizes were seen after two weeks (Figure 7). These patterns of cells were consistent with RBC production in the setting of mild hemolysis in a recovery process.

**Figure 6:**
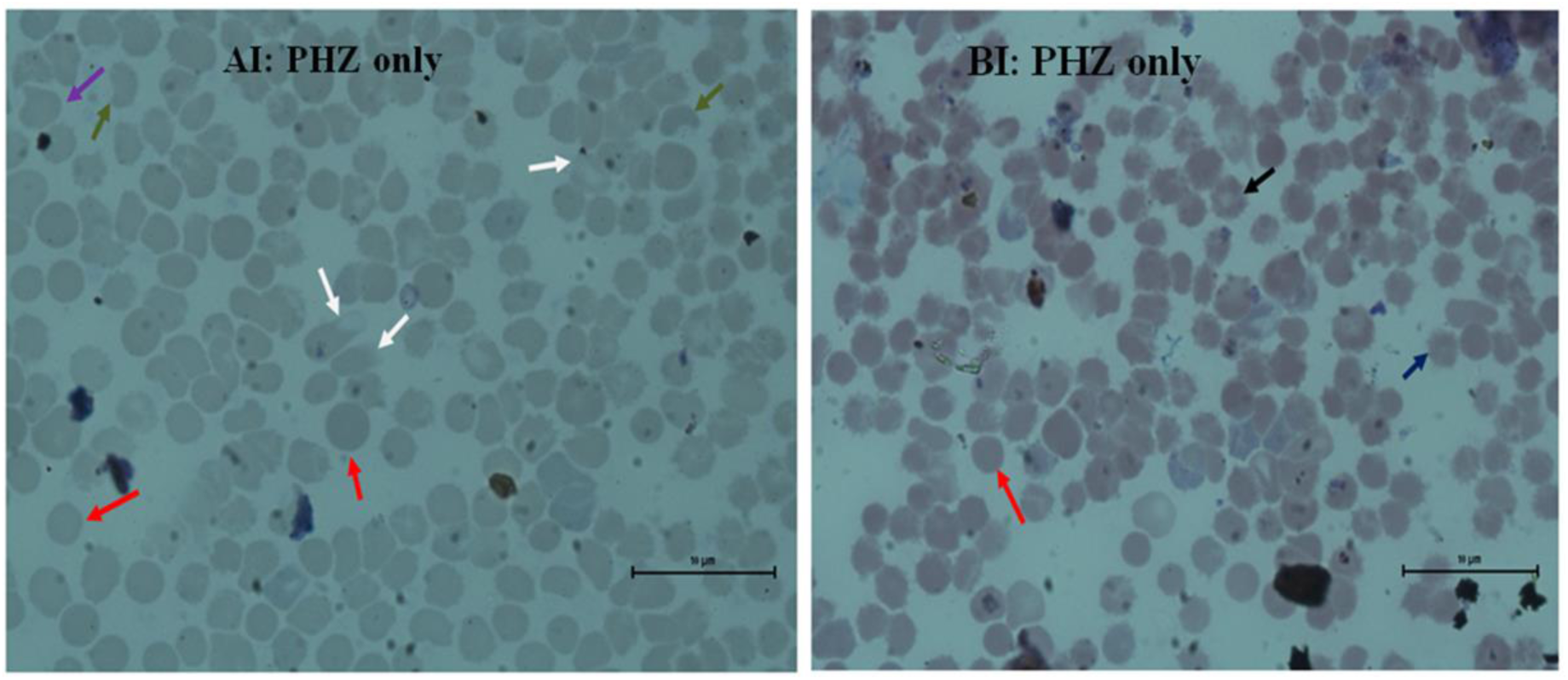
Blood smear, RBCs, Giemsa, x 100. **RBC morphological abnormalities.** Blister cells (bold white arrow), echinocytes/burr cells (bold black), tear drop cells (bold purple arrow), and acanthocytes (bold dark blue arrow). Tear drop cells are indicative mild marrow fibrosis (not diagnostic in this case). These patterns of cells were characteristic of an ongoing mild hemolysis secondary to oxidative stress (PHZ-induced).

**Figure 7:**
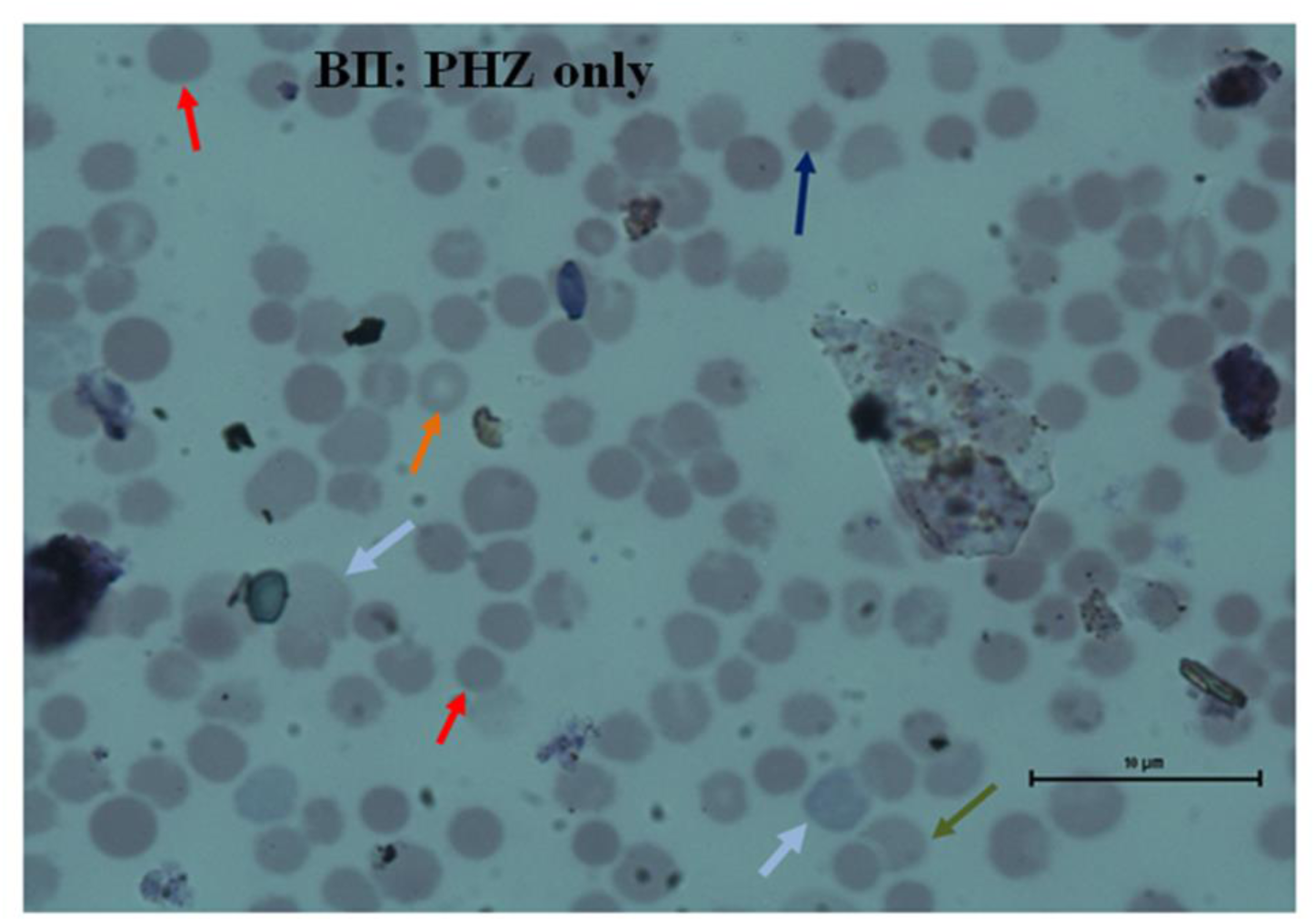
Blood smear, RBCs, Giemsa, x 100. **RBC morphological abnormalities**. Faint blue cells (bold light blue arrow), well hemoglobinised RBCs (bold dark orange), Bite cells (bold dark green arrow), acanthocytes (bold dark blue arrow), and spherocytes (bold dark red arrow). Faint blue cells are indicative of reticulocytosis.

#### Anemic treated with Ferro B syrup

The photomicrograph of these rats one week post therapy revealed moderately increased spherocytes of varied sizes, moderately increased normal sized & well hemoglobinised RBCs, slightly increased spur cells occasional tear drop cells, and few blister cells (Figure 8). After two weeks of treatment, the normal RBCs appeared more numerous compared to those that were treated for only one week. However, there were significant moderate populations of spherocytic cells (Figure 9). These morphological patterns of the RBCs that were seen in the smear of these rats were indicative of fairly adequate RBC production in the setting of mild hemolysis suggestive of relative recovery from the acute hemolytic episode induced by PHZ.

**Figure 8:**
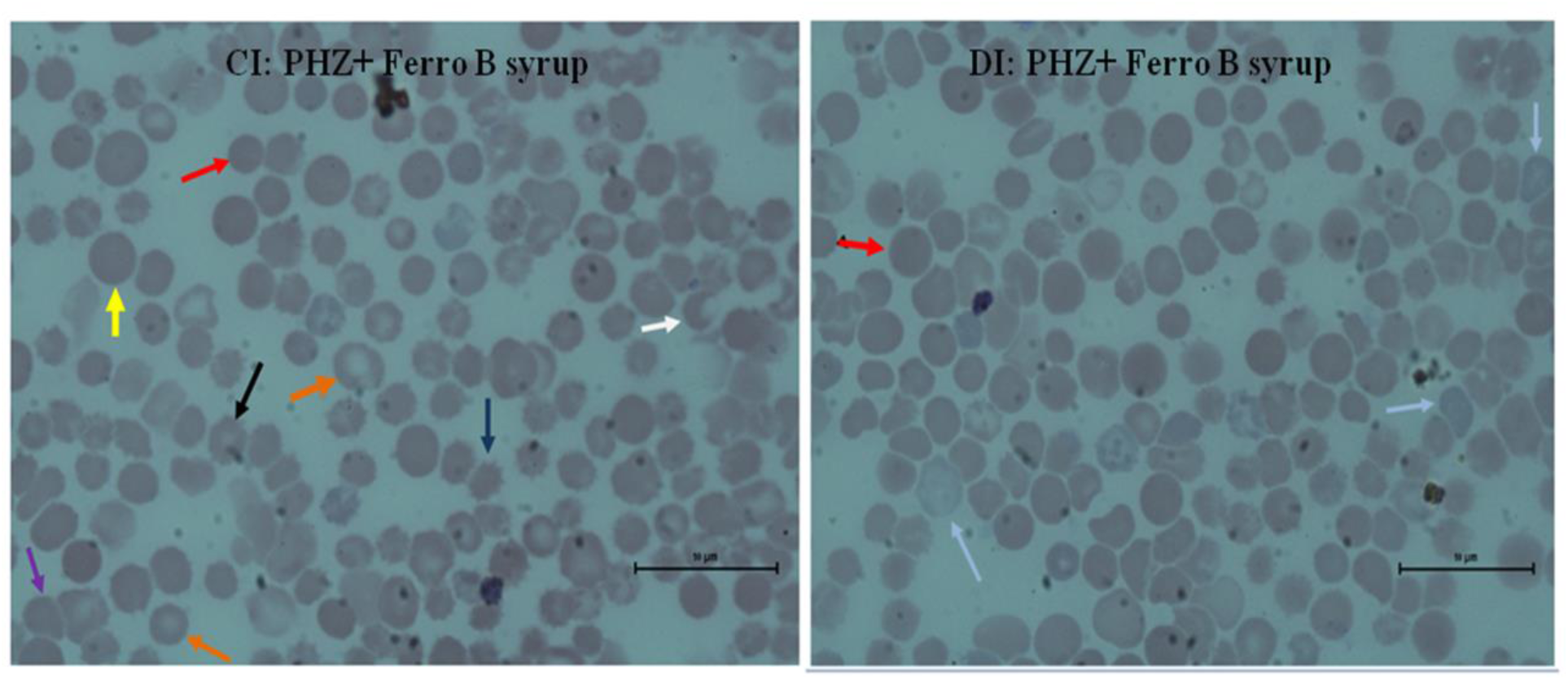
Blood smear, RBCs, Giemsa, x 100. Spherocytes (bold red and yellow arrows), RBCs (bold orange arrows), spur cells (bold dark blue); tear drop cells (bold purple arrow), blister cells (bold white arrow).

**Figure 9:**
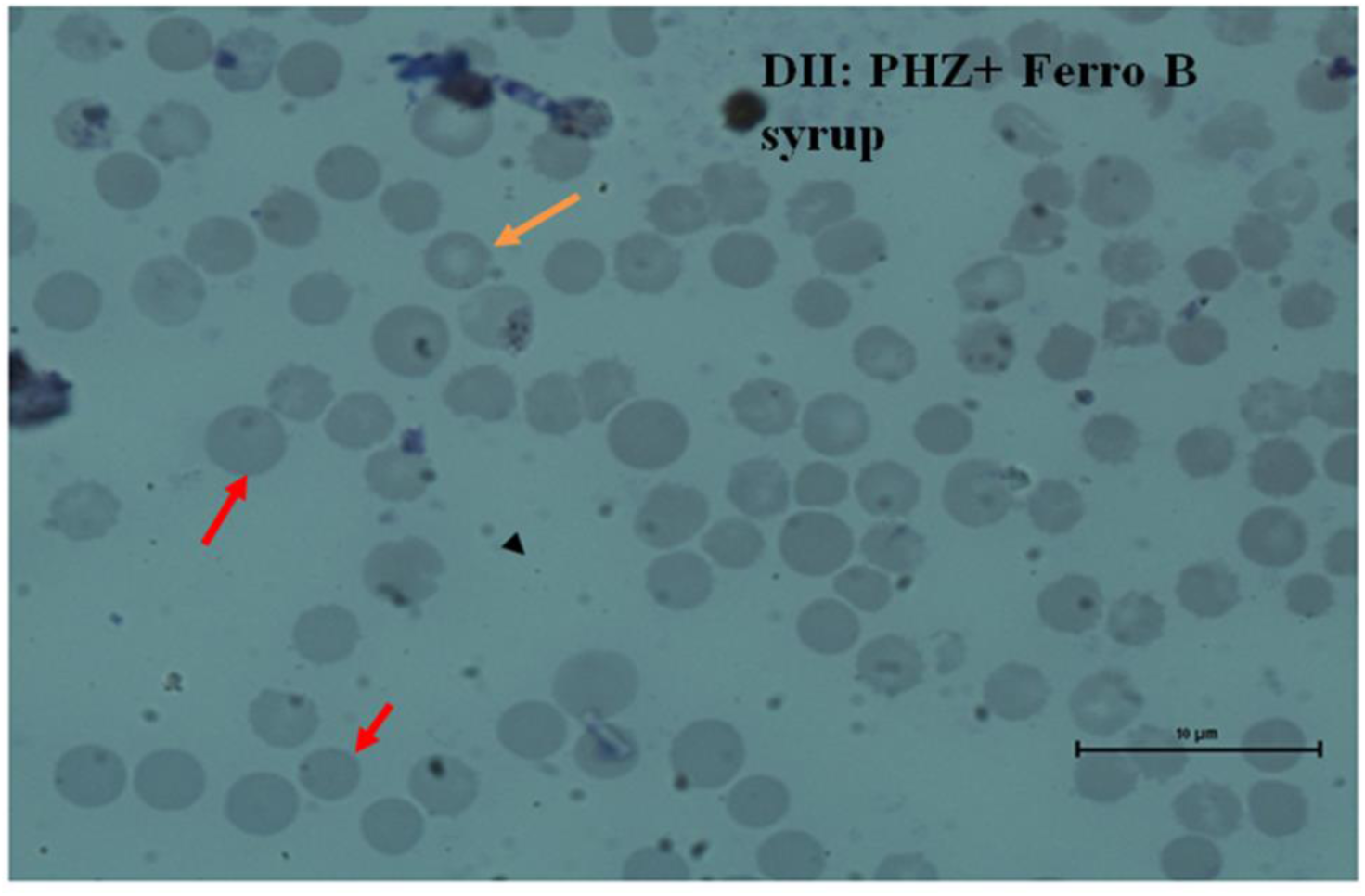
Blood smear, RBCs, Giemsa, x 100. Predominance of spherocytes & normal RBCs

#### Anemic treated with 400mg/kg of *T. indica* pulp extract

Moderate to numerous spherocytes, minor populations of spur cells/Acanthocytes, and a tear drop cell were seen. Few bite cells, blister cells, and fragmented RBCs were present. Moderately increased populations of normal RBCs were also apparent (Figure 10). The features seen were consistent with mild hemolysis and relative ongoing recovery in the course of the administered dose of the *T. indica* fruit pulp extract.

**Figure 10:**
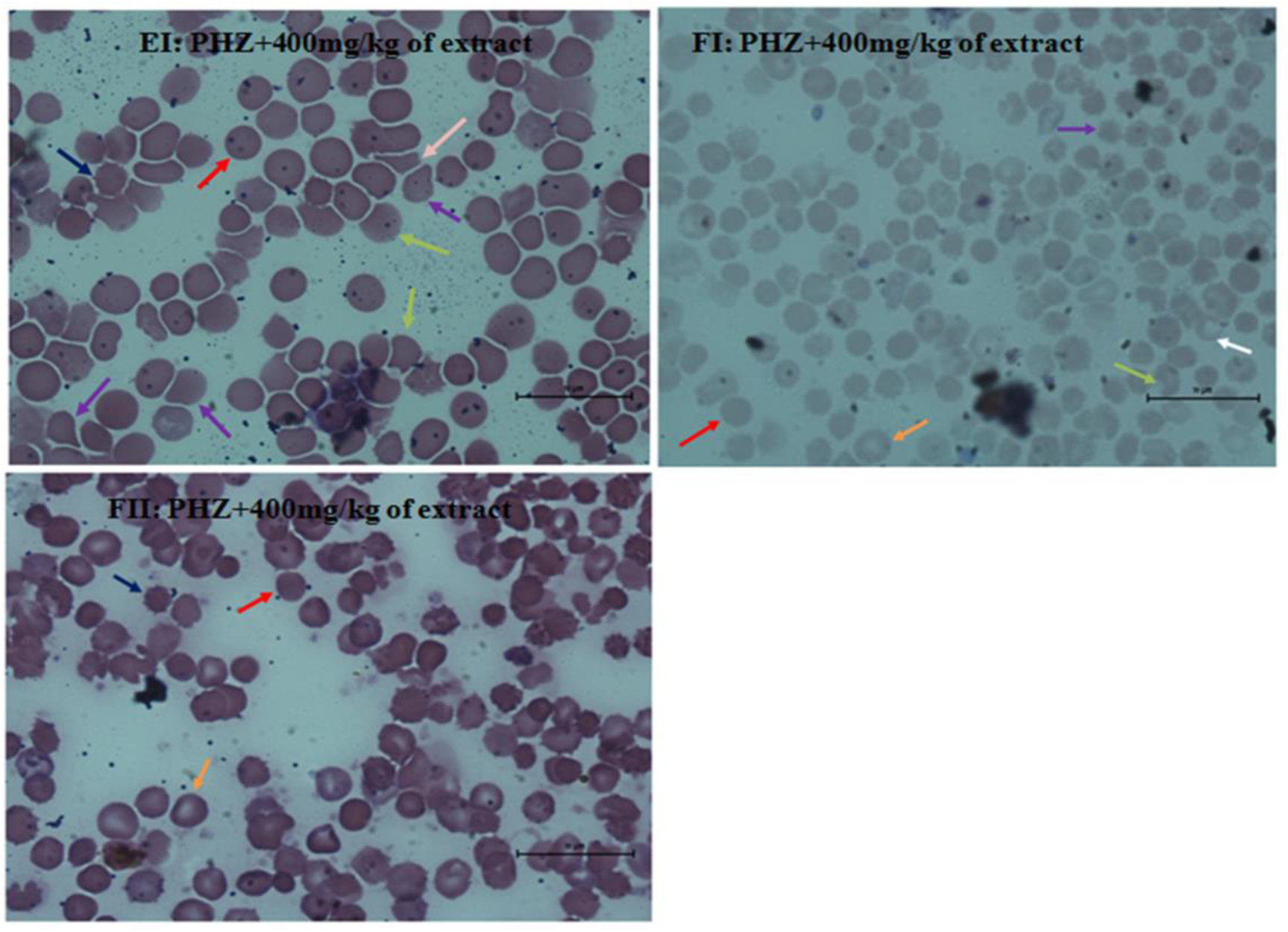
Blood smear, RBCs, Giemsa, x 100. Spherocytes (dense red arrow), Spur cells/Acanthocytes (Dark blue arrow), Tear drop cell (bold purple arrow), Bite cells (bold dark green arrow), Blister cells (bold white arrow), and RBCs (bold orange arrows).

#### Anemic treated with 800mg/kg of *T. indica* pulp extract

Moderate to large spherocyte population; few bite cells, blister cells, and tear drop cells were predominant. Spur cells were seen in the smear of the rats that were treated for one week (Figure 11). These pictures were typical with mild hemolysis and portrayed relative recovery following administered *T. indica* fruit pulp extract therapy at that dose (800mg/kg).

**Figure 11:**
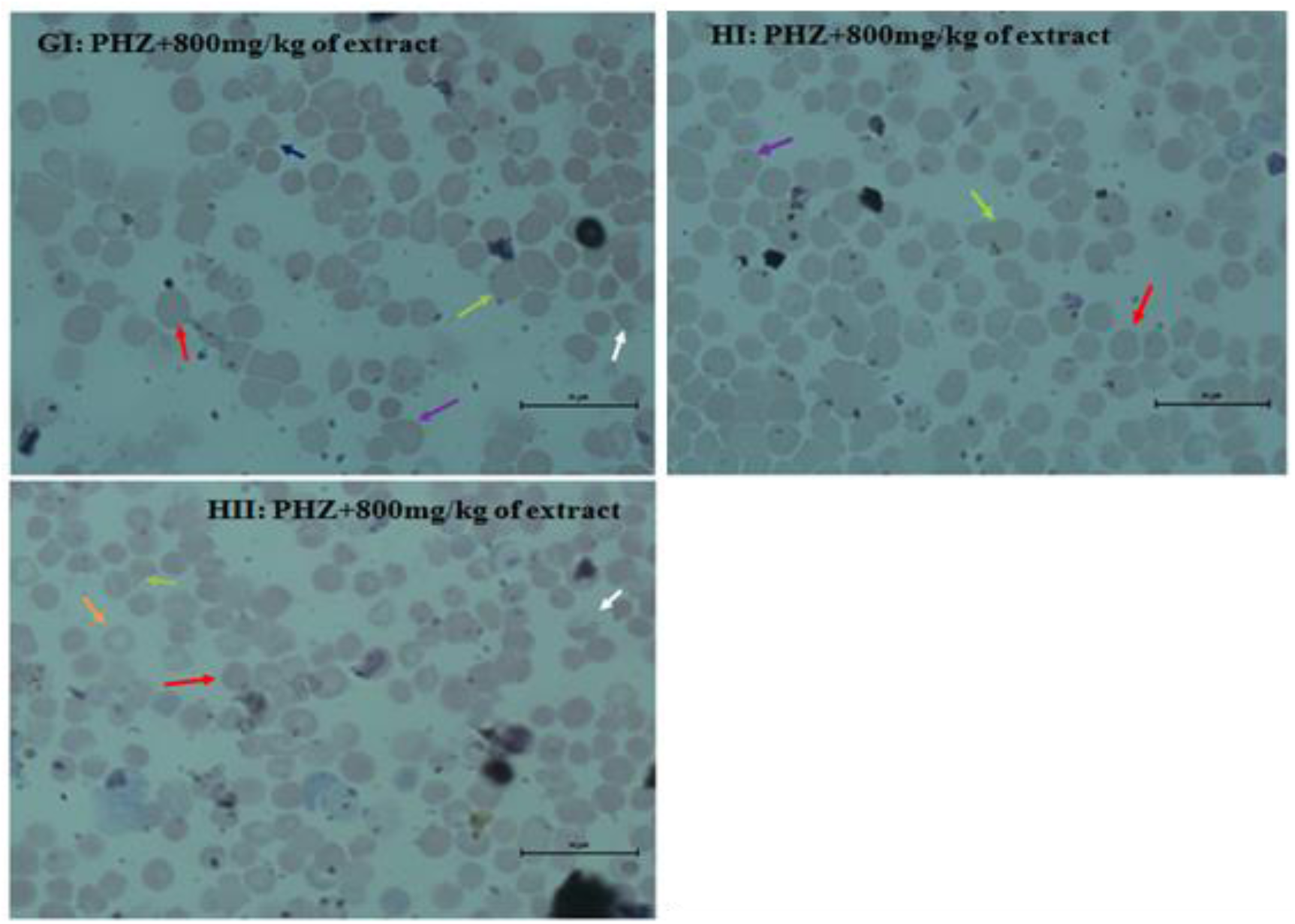
Blood smear, RBCs, Giemsa, x 100. Spherocytes (bold red arrow), Bite cells (bold dark green arrow), Blister cells (bold white arrow), Tear drop (bold purple arrow), and spur cells (bold dark blue arrow).

### Medium and high power bone marrow histomorphology results pre and post-therapy (extract, ferro B syrup and PHZ)

#### Medium power photomicrographs of Groups 1-5

The marrow cytoarchitecture were hypercellular in all cases, with the hematopoietic cells seen as prominent erythroid islands. Megakaryocytes (MK) were also present in all aspects. For the normal rats, the numbers of megakaryocytes seen were significant (about 16 cells per HPF) and showed normal characteristic morphology. However, in all the anemic rats (treated and untreated), the megakaryocytes displayed different characteristic morphology of abnormal nuclei patterns in terms of texture, number of lobes, color, separation, and nuclei: cytoplasm ratio. Vacuolated erythroid cells and adipocytes were seen in all the anemic rats (treated and untreated). Though Histiocytes were seen in the anemic rats that were receiving the extract, slight architectural distortion was seen in those that were receiving 400mg/kg of the extract (Figure 12).

**Figure 12:**
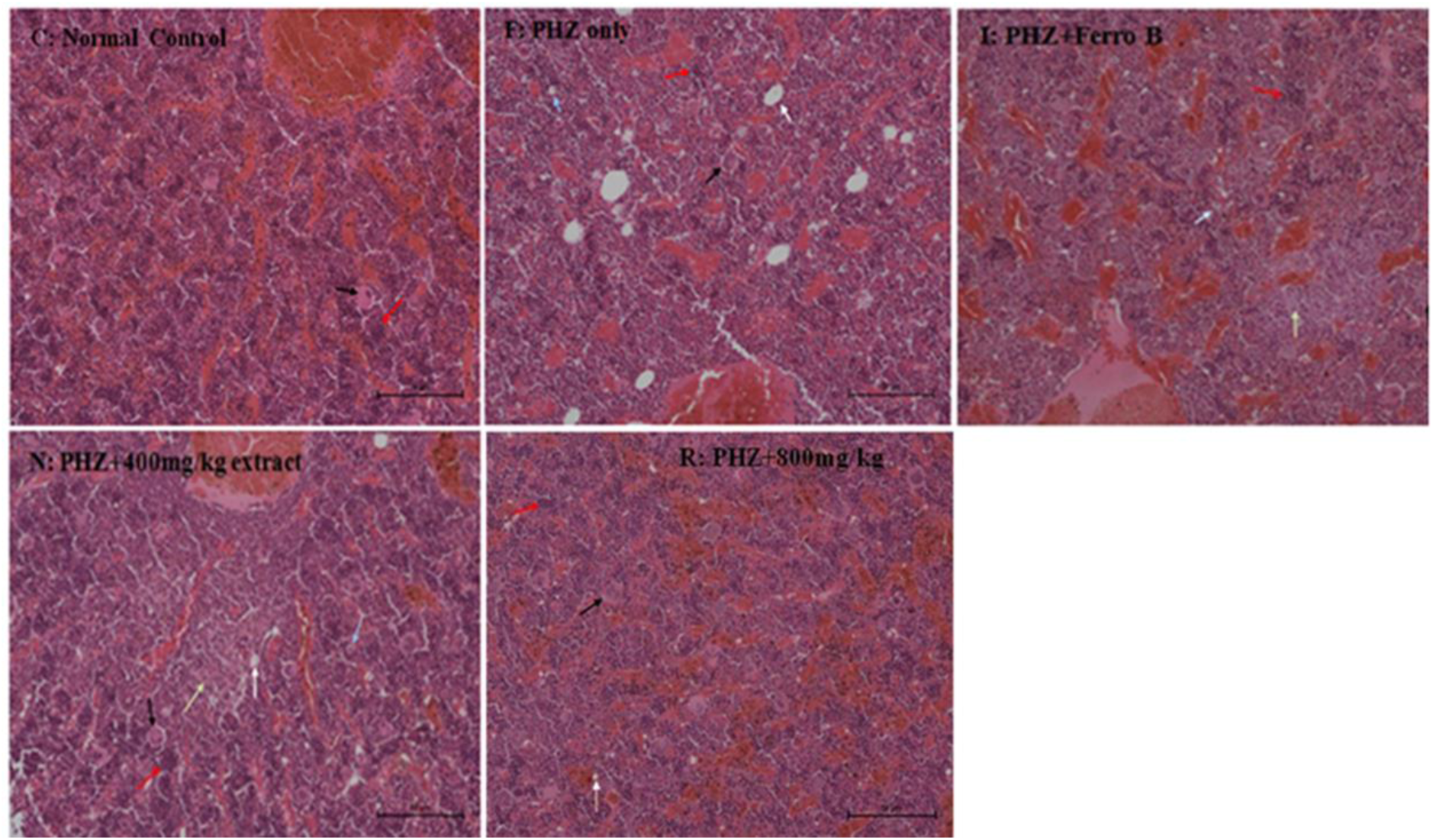
Hypercellular marrow, H & E, x20. Erythroid islands (red arrow), MK (black arrow), Adipocytes (white arrow), Vacuolated erythroid cells (light blue arrow), Histiocytes (light green arrow). **Panel C:** Normal marrow cytoarchitecture. **Panel F:** Adipocytosis; MK-distributed singly, nuclear lobes-hyperlobated & coarse. **Panel I:** Adipocytosis. **Panel N:** Adipocytosis, Focal Histiocytosis. **Panel R:** Adipocytosis.

#### High power photomicrographs of Groups 1-5

Though the cell types and cell characteristics in this magnification were not different from those at medium magnification, they appeared more visible and clearer. Areas of architectural distortion (swirling and lining of cells), the megakaryocytes, and the histiocytes were well defined (Figure 13).

**Figure 13:**
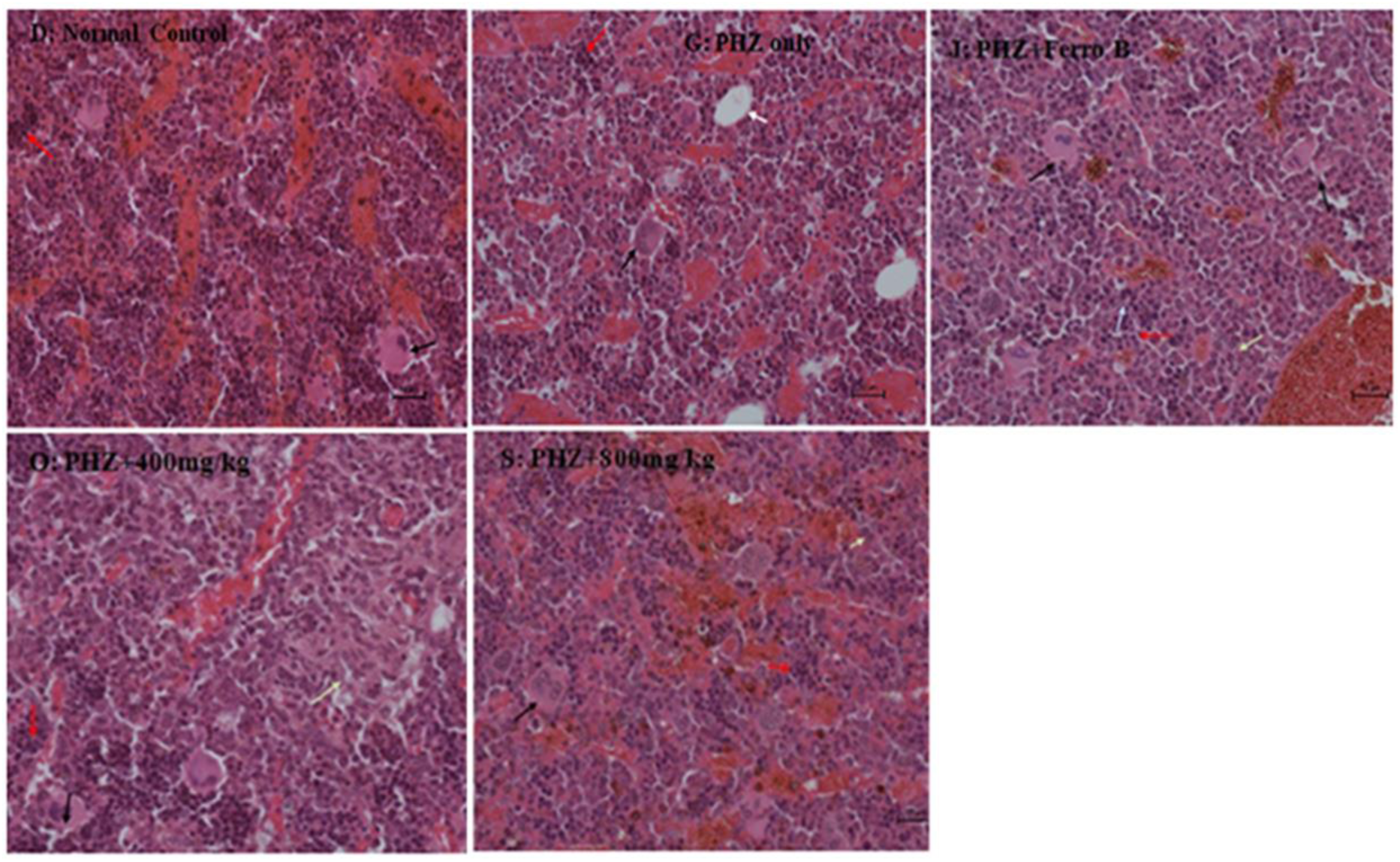
Hypercellular marrow, **H & E**, x40. Erythroid islands (red arrow), MK (black arrow), Adipocytes (white arrow), Vacuolated erythroid cells (light blue arrow), Histiocytes (light green arrow). **Panel D:** Normal marrow cytoarchitecture. **Panel G:** Adipocytosis; MK-dysplastic pattern. **Panel J:** MK are aberrant-separated nuclear lobes, hypolobation, and hyperchromatic. **Panel O:** Focal Histiocytosis; MK-abnormal morphology. **Panel S:** MK - dysplastic with abnormal patterns (large, coarse nuclear lobes are separated, nuclear: cytoplasm ratio is aberrant).

Histiocytosis (loss of marrow cells), adipocytosis, megakaryocytic dysplasia, and the vacuolated cells were degenerative features suggestive of PHZ-induced marrow fibrosis and marrow suppression. The significant erythroid hyperplasia was typical of an ongoing hematopoiesis indicative of active marrow regeneration following the therapies that were given.

## Discussion

### Effect of *T. indica* fruit pulp on serum LDH

In this study, PHZ was used as a model to induce hemolytic anemia; in particular DIHA, for the purposes of evaluating its influence on the therapeutic effectiveness of *T.indica* fruit pulp extract. The 60mg/kg body weight of the PHZ that was given intraperitoneally to the rat models daily for two days induced: marked thrombocytosis, generalized leucocytosis, low Hb level, microcytosis and macrocytosis, raised nucleated cells, and reticulocytosis; low MCH, MCHC and HCT, and elevated serum LDH. These differentials are typical in a hemolytic pattern, with accompanied paradoxical degree of myelofibrosis/marrow fibrosis, and may suggest a myeloproliferative disorder of a secondary cause, in this case PHZ. This agrees with the findings of Jaiswal *et al.*[33].

In this investigation, the biochemical levels of serum LDH was significantly raised in the anemic treated and untreated rats in the first and second week respectively, but more marked in the anemic-untreated, especially in the first week. This was due to marked RBC injury inflicted by PHZ’s various cytotoxic/or hemato-toxic mechanisms that resulted into its’ accelerated lysis and fragmentation [12, 34] intravascularly [35, 36, 3] and extravascularly within the splenic, hepatic, and bone marrow macrophages [36, 15] releasing massive amounts of the LDH enzyme into the plasma [1]. The observable 16% (58 IU/L) significant decrease extending to the second week was consistent with the resolution of hemolysis following therapeutic intervention. This was in agreement with Barcellini and Fattizzo [37], and Hillmen *et al*. [38]. A marked significant decrease in the serum LDH consistent with recovery from hemolysis was seen in the group receiving 800mg/kg bwt of the *T. indica* fruit pulp extract, in the second week. For those receiving 400mg/kg bwt of the extract and Ferro B syrup, their serum LDH were comparable. However, the group that didn’t receive the intended therapy, their serum LDH remained significantly high in the entire experimental period, due to persistent hemolysis.

While the serum LDH levels in the anemic-untreated rats remained unaltered (markedly high) throughout the experimental period, due to ongoing persistent hemolysis, 800mg/kg bwt of the *T.indica* fruit pulp extract caused a significant decrease in serum LDH activity in the second week consistent with significant recovery from hemolysis. The same extract, at a dose of 400mg/kg bwt, produced comparable serum LDH levels with Ferro B syrup. These results indicated that the intervention by the experimental molecule (*T. indica* fruit pulp extract) significantly decreased the serum LDH activity as reflected in the drop in the serum LDH levels. This could probably be due to *T. indica* fruit pulps’ physiologically active phytochemicals [39] and nutritional components [23] that: 1) have antioxidant properties, whose antioxidant molecules scavenged and neutralized the generated free radicals [40,17], decreased and prevented lipid peroxidation [41]; 2) modulated anti-oxidant enzyme activities that was involved in the reduction of oxidized proteins and lipids [42], deglutathionyled and reduced the protein disulphides formed [43]; 3) suppressed and modulated activities of immune cells and phagocytic cells, thus decreasing the production of autoantibodies by the B-cells and removal of antibodies from the RBCs [44]; T lymphocytes, CD4+T and CD8+T cell mediated cytokine generation [45] in response to antigens [46,47]; 4) restored lost electrolytes i.e. Sodium and Potassium, and minerals i.e. Calcium, Copper, Iron, Magnesium, Selenium and Zinc [23]. These possible mechanisms possibly corrected the defective/malfunctioned antioxidant system, restored electrolyte and mineral imbalances; and decreased autoantibody and drug dependent antibody generation induced by PHZ, reducing antibody mediated extravascular and complement mediated intravascular hemolysis [3]; thus conferring RBC membrane stability and RBC protection from PHZ induced hemolytic insult.

### Effect of *T. indica* fruit pulp on the reticulocytes

Reticulocytes, a highly sensitive marker in predicting erythroid changes [27], and monitoring recovery from hemolysis or “antihemolytic” therapy, are indicative of marrow hemopoietic activity in pathophysiologic states when present in the blood or seen in the peripheral smear [37].

In this study, PHZ induced acute hemolytic event in the rat models, evidenced by marked reticulocytosis, seen in the untreated group one week post-hemolysis induction is in line with Redondo *et al*. [48,49] who reported fulminant reticulocytosis 6-10 days post-PHZ administration. Though significant reticulocytosis was still observable after therapeutic intervention with ferro B syrup and the *T.indica* fruit pulp extract, mild reduction was only seen after 2 weeks of treatment in all the experimental groups. This observed pattern was likely due to: 1) stimulation of erythropoietin hormone following PHZ-induced hemolytic anemia that might have inhibited apoptosis of, and enabled terminal differentiation of the erythroid progenitor cells [50] leading to increased production of RBCs [51], at the same time decreased the number of autoantibodies present on each of the freshly formed RBCs from the aforementioned differentiation; 2) with concurrent suppressive and immune-modulatory effect of the *T.indica* fruit pulp extract on immune cells that led to decrease in the production of autoantibodies, and their removal from the RBCs [44].

### Effect of *T. indica* fruit pulp on the RBC morphology

PHZ induced gross RBC morphological distortions with spherocytic predominance, including bite cells and blister cells. Echinocytes, acanthocytes, tear drop cells, and faint blue cells were remarkably present. Spherocytes, the principal RBCs seen in immune mediated hemolytic anemia secondary to drugs or other causes, in this particular case, PHZ, might have resulted from antibody binding or complement that coated the RBC membrane, tagging it for destruction by the splenic macrophages (figure 6) [52], and/or osmotic lysis [30]. Whereas bite cells and blister cells were characteristic of oxidative damage and unstable Hb; the faint blue cells and tear drop cells were indicative of reticulocytosis and myelofibrosis i.e. secondary (PHZ-induced) marrow fibrosis (figure 12 & 13) [52].

Though the morphological patterns of abnormal RBCs in the anemic treated rats were comparable, normal RBC populations observed after 2 weeks ranged from few in the untreated anemic rats to fairly adequate in the anemic rats receiving Ferro B syrup, and moderately increased in the anemic rats receiving the respective doses of the *T.indica* pulp extract, significant of relative recovery from hemolysis. This was likely due to: a) compensatory bone marrow erythropoiesis [2] demonstrated by the few normal RBCs in the anemic untreated rats; and b) combination of both compensatory erythropoiesis [2] and stimulated erythropoiesis [30] following treatment, in response to PHZ-induced hemolytic effect.

### Effect of *T. indica* fruit pulp on the bone marrow histomorphology

According to this investigation, PHZ induced bone marrow fibrosis, marrow suppression, and other degenerative changes. The marrow fibrosis was characterized by megakaryocytic dysplasia seen as dysplastic and abnormal patterns coupled with focal histiocytosis. Marrow suppression and other degenerative changes were featured by multifocal adipocytosis and the presence of vacuolated erythroid cells. Adipocytosis was more marked in the anemic untreated rats compared to the anemic treated that showed scanty adipocytes that were negligible, following therapy.

Though PHZ caused marrow cyto-architectural distortion, it paradoxically increased the number of erythrocyte-committed progenitors and colony-forming units [53] seen as erythroid islands/clusters in the bone marrow, and reticulocytosis in the peripheral smear. Following the test therapeutic intervention, marrow regenerative features were typical, marked by marrow hypercellularity characteristic of erythroid hyperplasia, a principal feature suggestive of on-going hematopoiesis, and markedly decreased adipocytes. These regenerative changes were most likely due to the *T. indica* fruit pulp extract whose phytochemicals and micronutrient components might have: 1) enhanced the immune cells by modulating the cell mediated immune response thus balancing their suppressive activities and cytokine generation; 2) enhanced the marrow microenvironment (stroma and matrix) that recognized and retained Hematopoietic Stem Cell [54], and provided the hematopoietic growth factors required for progenitor cell proliferation, differentiation, and maturation [55]; 3) provided the extracellular requirements for nucleic acid synthesis [56], and precursor molecules e.g. Fe and proteins required by the erythroid precursors to synthesize Hb [57]; 4) Modulated lipid metabolism associated genes [58,59] that resulted in marked decrease in marrow adipocytes.

## Conclusion

*T.indica* fruit pulp extract effectively stimulated hematopoiesis in response to drug induced hemolytic effect on the hematopathologic parameters, with significant improvement from hemolytic anemia, as evidenced by the marrow regeneration, significant reduction in serum LDH activities, peripheral reticulocyte circulation, and the presence of significant populations of normal mature RBCs in the peripheral circulation within 14 days. This was achieved at the doses of 400mg/kg & 800mg/kg of the extract. Physiologically active phytochemicals and micronutrient elements contained in the *Tamarind* pulp are most likely components accounting for its remarkable hematopoietic potential. However, the acanthocytes, echinocytes, and teardrop cells are suggestive of hepato-renal involvement.

## Acknowledgement

The authors are grateful to Mr. Alinaitwe Lordrick for his help in tissue processing, Mr. Sengendo Ibrahim for the thin Blood smear preparation and reticulocyte determination, Mr. Elias Kwizera for LDH analysis. Much appreciation goes to Dr. Eneku Wilfred, pathologist, for reviewing the histopathology slides.

## Author Contributions

**Conceptualization:** Cherop Patrick Stephen.

**Data curation:** Cherop Patrick Stephen, Dare Samuel Sunday.

**Formal analysis:** Cherop Patrick Stephen, Dare Samuel Sunday.

**Investigation:** Cherop Patrick Stephen, Edgar Fernandez Mario, Dare Samuel Sunday.

**Methodology:** Cherop Patrick Stephen, Dare Samuel Sunday.

**Project administration:** Cherop Patrick Stephen, Edgar Fernandez Mario, Dare Samuel Sunday.

**Resources:** Cherop Patrick Stephen, Dare Samuel Sunday.

**Supervision:** Edgar Fernandez Mario, Dare Samuel Sunday.

**Writing – original draft:** Cherop Patrick Stephen, Dare Samuel Sunday.

**Writing – review & editing:** Cherop Patrick Stephen, Dare Samuel Sunday.

## Notes

### Competing Interest Statement

The authors have declared no competing interest.

